# T cell repertoire diversity measurement; inferences from a dynamical systems model, Fourier Analysis of the T cell repertoire

**DOI:** 10.64898/2026.08.24.746887

**Authors:** Amir A. Toor, Alejandro Marinos-Velarde, Rehan Qayyum

## Abstract

T cell repertoire sequencing has unveiled a vast, complex array of T cells responsible for the human immune responses. Traditional analytic methodology fails to fully characterize and quantify the diversity of T cell receptors constituting the T cell repertoire. T cell receptor clonal frequency measured in terms of T cell receptor beta (TRB) V gene segment usage when arrayed in correspondence with the respective V gene segment positions on the TRB loci yields a periodic, undulating curve in the spatial domain of the TRB genomic locus.

Using the genomic distance from the TRB-D1 segment to the TRB-V1-29 segments, Fourier analysis was performed utilizing Lomb-Scargle periodogram to obtain Spectral Power curves quantifying the TRB V clonal frequencies from 6 allogeneic stem cell transplant donors (baseline) and recipients (≥100 days) using a variety of analytic software. Spectral Power curves revealed dominant spectral peaks at wavelengths ranging from 4-9 kb (113-252 millicycles/kb) in the six donors, with consistent frequency domain spectral patterns. This is consistent with similar use of V segments across healthy individuals. Recipients on the other hand demonstrated more dispersed spectra, with a spectral centroid shifted towards higher frequencies compared to donors (260 vs. 247 millicycles/kb). Consistent with this observation, the Low Frequency Index was lower in the recipients (0.18 vs 0.20). Power was concentrated in the <3 kb and 3-12 kb wavelengths in both groups. The analyses reported here demonstrate that the healthy SCT donors have a remarkably similar spectral signature occupying short to intermediate wavelegnths in the frequency domain, whereas recipients tend to shift towards higher frequencies. These findings are consistent with a normal organized distribution of TRB V segment usage in healthy individuals (by analogy other loci), and a more diffuse and disorderly usage in recipients, consistent with the notion of T cell responses constituting a dynamical system which evolves as a function of time. Fourier analysis of TRB (and potentially TRA) sequencing data provides a repertoire wide summary of T cell clonal distribution.

## Introduction

Sequencing the T cell receptors using next generation sequencing (NGS) platforms yields an immense amount of data for samples evaluated, with millions of unique T cell receptor (TCR) alpha or beta sequences identified, along with their frequency. The complexity of these data sets limits their utility in direct clinical practice, making TCR sequencing a research tool where these data are often summarized using analytic methodology from information theory. Such concepts as Shannon’s entropy are utilized to represent large data sets in a compact fashion and subject TCR data from different patients to statistical analysis and draw appropriate inferences regarding the clinical impact of T cell repertoire recovery. Nevertheless, these indices yield values which are not truly reflective of the underlying biological complexity, and result in loss of information, rendering the ensuing correlations approximate at best. Therefore, alternative analytic methodology is needed to summarize TCR repertoire data, which may more accurately reflect underlying biology and provide an intuitive, quantitative understanding of T cell repertoire evolution over time in different clinical scenarios.

The human T cell repertoire has several organizational principles which have been previously elucidated and may be utilized in developing new methods for studying repertoire diversity. First, TCR defined clonal frequencies are logarithmically scaled and have Power Law distributions. Inherent to this quality, when clonal frequencies are assessed from a TCR-beta (TRB), Variable, Joining and Diversity (VDJ) segment defined clonal frequency perspective, these have a fractal ordering, i.e., distribution of T cell clones remains proportional across different levels of V,D and J segment recombination .^1^ Secondly, when the T cell receptor V, D and J segment locations are examined on the TCR loci, these are highly ordered, with segment lengths and the inter-segment distances (in base pairs) being logarithmically organized. V gene segments are the longest, J gene segments have an intermediate length and D gene segments are the shortest. Finally, when the TCR beta defined clonal frequencies are examined as a function of the V and J segment positions on the TCR beta locus on chromosome 7q,^2^ a periodic distribution of clonal frequencies is observed as the clonal frequencies are mapped across the TCR beta locus from the 5’ to the 3’ end of the locus.^3^ This spatially determined periodicity in the likelihood of TCR beta V and J segment recombination has also been observed in TCR alpha defined clonal frequencies (*manuscript in preparation*). Given this mathematical organization of the repertoire and that immune responses following stem cell transplantation (SCT) behave as dynamical systems,^4, 5^ logically the T cell repertoire should behave in a broadly quantifiable manner. Such broad characterization of the T cell repertoire may permit a deeper understanding of large-scale phenomenon observed post-transplant.

Fourier analysis is a method of analyzing periodic phenomenon, such as waves travelling across different media (e.g., soundwaves propagating through air). The underlying assumption is that complex waveforms (such as a multi-instrument melody) represent a sum of many simple sine waves (fundamental harmonics) which combine to yield the final note. In other words, several waves of different frequency (number of oscillations over unit time) of a certain amplitude (magnitude) combine to yield a complex wave. ECG or EEG patterns are similar periodic phenomenon with complex waveforms. Fourier analysis is a mathematical method which allows the complex waves to be deconstructed into constituent simple harmonic waves of different frequencies, and calculates the contribution of each unique frequency to the overall amplitude of the combined wave. This is accomplished by a mathematical procedure called Discrete Fourier Transformation (DFT). The DFT takes the amplitude (height) of a wave from the time domain (amplitude plotted on the y axis, as a function of time on the x axis), and calculates the contribution of each constituent frequency comprising the complex wave: in turn plotting the amplitude in the frequency-domain (the magnitude contribution of each frequency to the final amplitude of the complex wave, Power). The relevant frequencies comprise the spectrum encompassed by the complex waves. As an example (Figure X) two simple waves (Blue: low frequency/long wavelength, Red: High frequency/short wavelength) combine to form a complex wave (Purple combined bimodal wave), which when transformed and visualized in the frequency domain can be seen to be comprised of two waves of different frequencies (thus different wavelengths) contributing different amplitudes to the final wave. This comprises the frequency spectral representation of the complex wave.

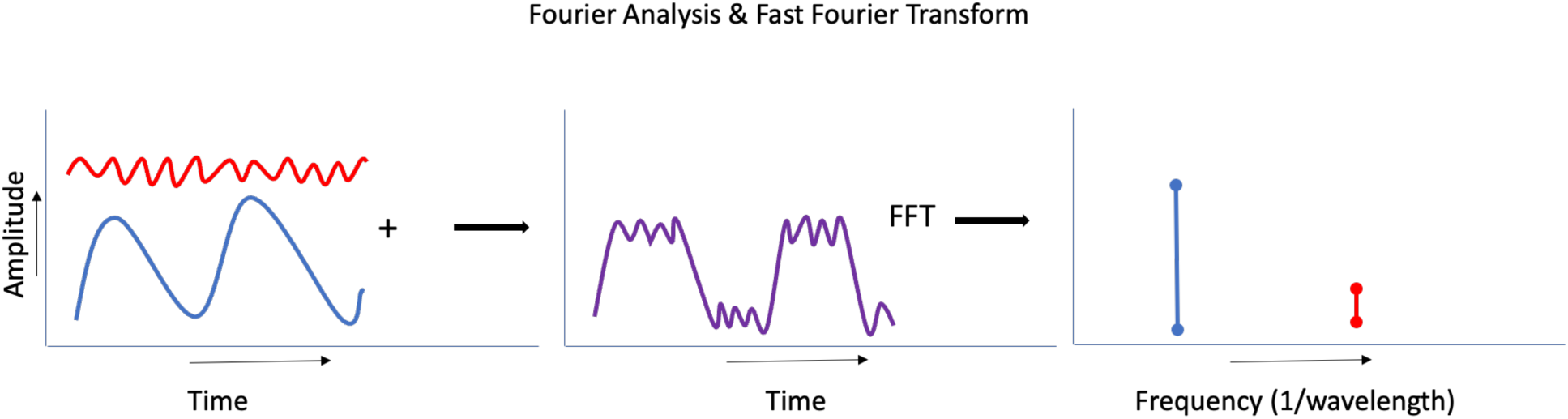

This concept of oscillation across the time domain to produce a periodic phenomenon (wave) may be similarly extrapolated to any cyclic variation in phenomenon with a measurable output, such as the probability of VDJ recombination occurring across a strand of double helical DNA molecules with V and J genes sequentially organized on it; in this instance the periodic phenomenon of interest is variation in the probability of TRB V and J genes participating in recombination and contributing to the final repertoire (represented by T cell clonal frequencies), producing oscillations mapped to a spatial domain, such as the TRB locus (measured in base-pairs of DNA). Similar to time domain-based phenomena (sound waves), this spatial waveform quantifying the sequential V gene segment defined T cell clones, may also be examined in the frequency domain to estimate the relative contribution of V gene segments to the final T cell clonal repertoire. Therefore, when the T cell repertoire at different time points is measured, any change in the relative TRB (or TRA) gene segment usage may be summarized and information regarding T cell clonal evolution be summarized, with greater preservation of information. Fourier analysis does so by decomposing the final T cell repertoire to highlight the relative contribution of different regions of the TRB gene. This method may prove to be a versatile tool for comparing T cell repertoire both within and between individuals and over time. ^6^

## Methods

Anonymized data from patients who underwent allogeneic peripheral blood stem cell transplantation from HLA matched unrelated donors on an IRB approved protocol at the Virginia Commonwealth University was utilized. SCT donor and recipient samples for determining T-cell clonal frequency were obtained as part of a clinical trial approved by the institutional review board at Virginia Commonwealth University (ClinicalTrials.gov Identifier: NCT00709592). T cell repertoire evaluation was performed on the donor product, and the recipients at ≥ day 100 or one-year post transplant, using Adaptive Biotechnologies’ T cell receptor Clonoseq assay (Adaptive Biotechnologies, Seattle, WA).^1, 3^ TRB V sequence reads (corresponding to T cell clones) were tabulated sequentially according to the 5’ to 3’ order of their V gene segments on the TCR beta (TRB) locus, along with the genomic distance of the respective locus from the TRB D1 gene segment. A frequency of zero was given to the TRB V pseudo genes. The spatial dimension of the TRB V gene segments was the angular distance (in radians) from the TRB D1 segment;^3^ approximated by KB for ease of understanding. Of note, TRB V30 is located 3’ of the TRB D gene segment so was excluded from the reported analyses. Data were obtained from six transplant donor recipient pairs (*previously published in Toor et al, 2016*).^3^ All the patients presented here underwent HLA MUD SCT (8/8, n=5; 7/8, n=1 – DRP 14/HLA B mm). Patients 5, 8, 9 and 14 developed GVHD during the study period/sample acquisition time. Samples 3 and 5 were collected at day 100 (R2) and the remaining samples between days 100 - 365 (R3).

### Spectral Analysis Methodology

Spectral analysis was performed using a custom Stata pipeline (StataNow 19, StataCorp, College Station, TX) supplemented with Python 3 (scipy.signal.lombscargle, numpy, pandas, matplotlib) for numerical computation and visualization. The analysis follows the Lomb-Scargle periodogram approach applied to raw, unevenly spaced V-gene clonal frequency data, avoiding the interpolation artifacts that can arise with conventional Fast Fourier Transform (FFT) methods.

For each sample (12 total: 6 donors and 6 recipients), the TRB V gene segment defined clonal frequencies (copy number) were arrayed according to the genomic position of the respective gene segments on chromosome 7q, measured in radians (calculated form kilobases (kb) of DNA base pairs) from the TRB D1 gene segment. The genomic coordinates of the V gene segments are inherently unevenly spaced, reflecting the physical organization of the TRB locus. The Lomb-Scargle periodogram computes spectral power at each candidate frequency by performing a least-squares fit of a sinusoidal model to the raw data, making it suitable for unevenly sampled spatial data without requiring interpolation.

The frequency grid was defined from a minimum frequency f_min_ = 1/60 kb (0.016 cycles/kb [16 millicycles/kb]; corresponding to a maximum wavelength of 60 kb) to a maximum frequency f_max_ = 0.5 cycles/kb [500 millicycles/kb] (0.5 cycles/kb; wavelength 2 kb; Nyquist limit for the minimum inter-V-gene spacing). The frequency grid consisted of 1024 evenly spaced bins on the linear frequency scale. The relevant section of the TRB locus spanned, between TRB-V1 and TRB-D1 ∼550 kb. The standardized frequency window ensured reproducibility across analyses by fixing the long-wavelength cutoff at 60 kb, rather than using a data-dependent cutoff such as 1/(genomic span), which varies with the number of V genes included. An important consideration here is that unlike time dependent periodic phenomenon, the TRB clonal frequency waveform has no negative component, with all numeric values going from 0 to *x*. It is also important to recognize that the frequency spectrum represents a compilation of calculated (virtual) spatial frequency bands which summed together yield the final TRB V gene segment defined T cell clonal frequency distribution across the locus. These bands will therefore represent the relative contribution of all the V gene segments across than the entire TCR locus.

### Spectral Metrics

The following metrics were derived from each power spectrum. The spectral centroid (SC) is the weighted mean frequency of the power spectrum, calculated as Sum (f_i * P_i) / Sum(P_i), where f_i is the frequency and P_i is the power (amplitude, or T cell clonal copy numbers) at that frequency. In essence the Fourier analysis determine the frequency spectrum which best represents the repertoire; SC captures the central tendency of the spectral distribution. Higher SC values indicate power concentrated at shorter wavelengths (higher frequencies). This would imply relative flattening of the TRB repertoire periodicity, or a diminution of the periodicity observed, with more uniform contribution of TRB V gene segments. The low-frequency index (LFI) is the fraction of total spectral power below 100 millicycles/kb (10 kb wavelength). Spectral entropy is the Shannon entropy of the normalized power distribution, measuring the uniformity of power spread across the frequency grid, with higher values corresponding with greater randomness across the frequency spectrum, corresponding to a more disordered repertoire in terms of V gene segment contributions. Band power is the fraction of total spectral power within defined wavelength intervals: 3-12 kb (Short Spacing), 12-30 kb (Intermediate Spacing), 30-60 kb (Long spacing), and (Ultra-long pacing) >60 kb. The residual band (<3 kb) captures power at the highest frequencies. In essence this indicates the virtual frequencies which contribute to the greatest extent in the final T cell repertoire generation, and whether these follow established periodicity or vice versa.

Analysis was repeated using MATLAB R2026a (The MathWorks, 2026) with the Signal Processing Toolbox add-on. Spectral analysis was performed using the Lomb-Scargle algorithm to identify periodicities in the data.

Fast Fourier Transform was not used since genes are not evenly spaced in the genome. Signal intensities for both patient and donor datasets were standardized using z-score normalization (mean = 0, standard deviation = 1) prior to analysis to control for amplitude differences between samples. The reported parameters were spectral centroid, and Shannon’s entropy.

### Jensen-Shannon Divergence

The Jensen-Shannon Divergence (JSD) was computed between each donor-recipient pair to quantify the overall dissimilarity of their normalized power spectra. JSD is a symmetrized version of the Kullback-Leibler divergence, bounded between 0 (identical spectra) and ln(2) (maximally different). Unlike single-metric comparisons (e.g., SC or band power), JSD captures differences across all frequency bins simultaneously. Both natural and binary (log2) units are reported.

### Bootstrap Confidence Intervals

To quantify uncertainty arising from the limited number of V gene observations (n = 67), non-parametric bootstrap resampling was performed. For each sample, 1000 bootstrap replicates were generated by resampling the 67 V gene observations with replacement. The Lomb-Scargle periodogram and all spectral metrics were recomputed for each replicate. The 95% confidence interval was taken as the 2.5th and 97.5th percentiles of the bootstrap distribution, reported alongside the median.

### Sensitivity Analysis

The sensitivity of the band power metric was assessed by varying two key analytical parameters: the number of frequency bins (128, 512, or 1024) and the minimum frequency cutoff, f_min_ (1/span, 1/120 kb, or 1/60 kb). Six combinations were evaluated for each pair. This analysis establishes the robustness of band-level summary measures to grid resolution and frequency window choice.

### Supplementary Sensitivity Analyses

Two supplementary analyses were performed to verify that specific data features do not artifactually drive the spectral results. First, a pseudogene comb-filter test excluded all 18 TRB V pseudogenes (retaining 49 functional V genes) and recomputed all metrics. This test addresses the concern that zero-frequency data points may introduce a periodic artifact into the power spectrum. Second, a V30 exclusion sensitivity analysis tested whether the inclusion of TRB V30 — which lies 3’ of the TRB D1 segment at position -12.6 kb and was excluded from the original manuscript’s primary analysis — meaningfully alters the spectral metrics.

The complete analysis pipeline is implemented in the do file fft_dr_reproduction.do, which is available as supplementary material. The full Stata log and all output tables are provided in the supplement.

Initially, Fourier analysis was performed on these organized TRB V clonal frequency data from the 6 DRP, using Claude (claude.ai). Because of the unevenness (biological variance) of V gene locations on the TRB locus, a Lomb-Scargle correction was applied by Claude.ai at the time of analysis. Supplemental analyses were carried out using ChatGPT 4.0 (chatgpt.com) to verify the trends observed in Claude. Following initial proof of concept analyses, Fourier analysis was performed as detailed below.

## Results

### Overview of Spectral Analysis

The TRB V clonal frequency distribution in the spatial domain exhibited a periodic oscillatory pattern in all samples, as previously reported.^3^ To obtain the frequency domain representation of these data, Lomb-Scargle periodogram analysis was performed on TRB V gene clonal frequency data from 6 donor-recipient pairs (DRPs), yielding 12 power spectra sampled on a 1024-bin frequency grid with a standardized frequency window (f_min_ = 1/60 cycles/kb, f_max_ = 0.5 cycles/kb). Decomposition of this spatial waveform into its constituent frequencies via the Lomb-Scargle periodogram revealed dominant spectral peaks at wavelengths of approximately 4-9 kb (113- 252 millicycles/kb) in all six donors, corresponding to the spacing between V gene clusters on the TRB locus. The most striking feature was the uniformity observed in the spectra of donors, consistent with a similar usage of TRB V gene segments across the normal donors studied (**Figure 1**). This was in distinction to the recipient spectra which were considerably more random in the distribution of their spectra (**Figure 2**). Individual DRP comparison demonstrated a small, albeit consistent shift in spectral frequencies in the recipients (**Supplementary Figure 1**). The Power spectra depicted as a function of wavelengths also demonstrated a similar organized state in donors transitioning to disorganization in recipients (**Figures 1 & 2**).

**Figure 1.**
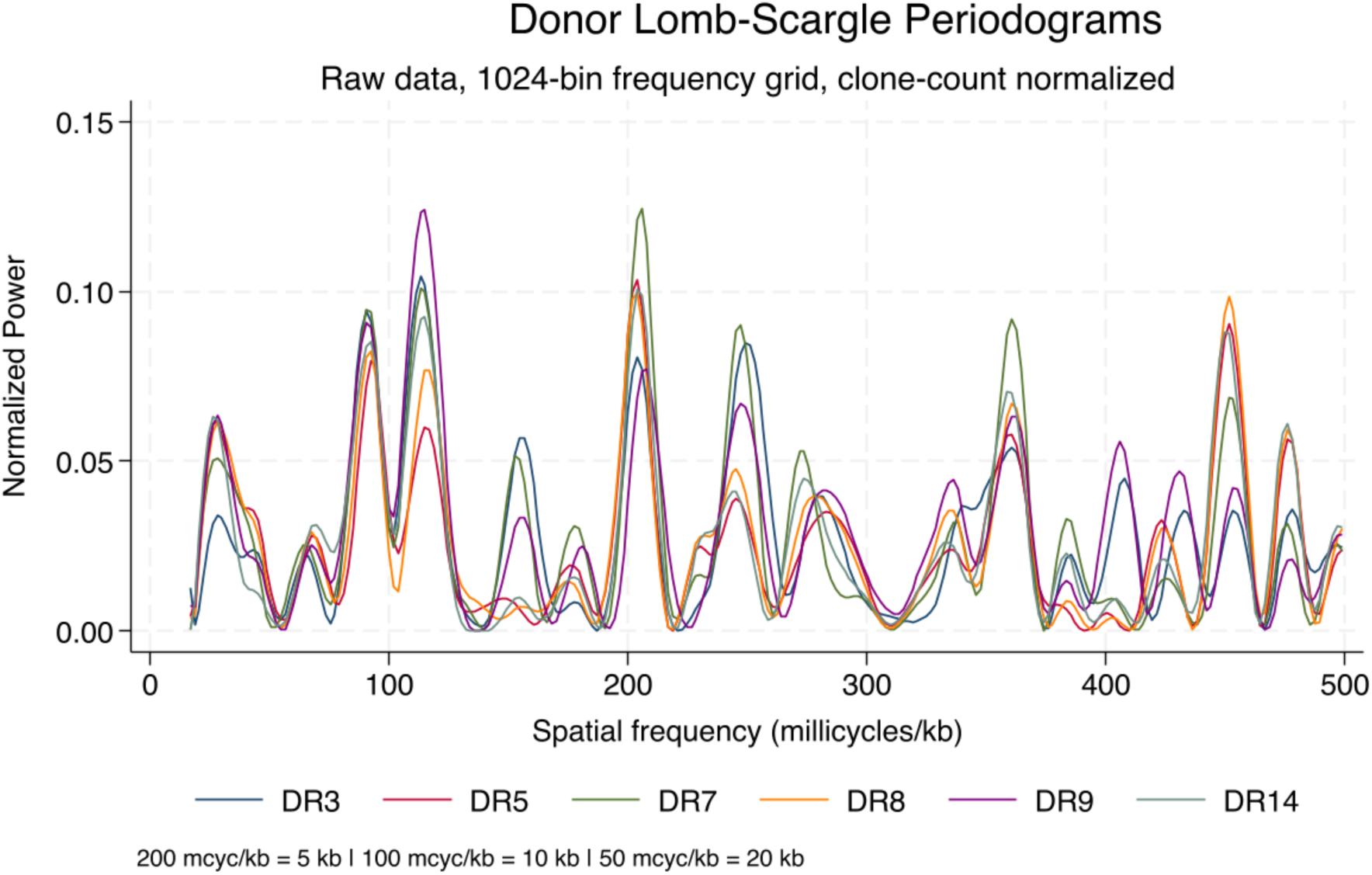
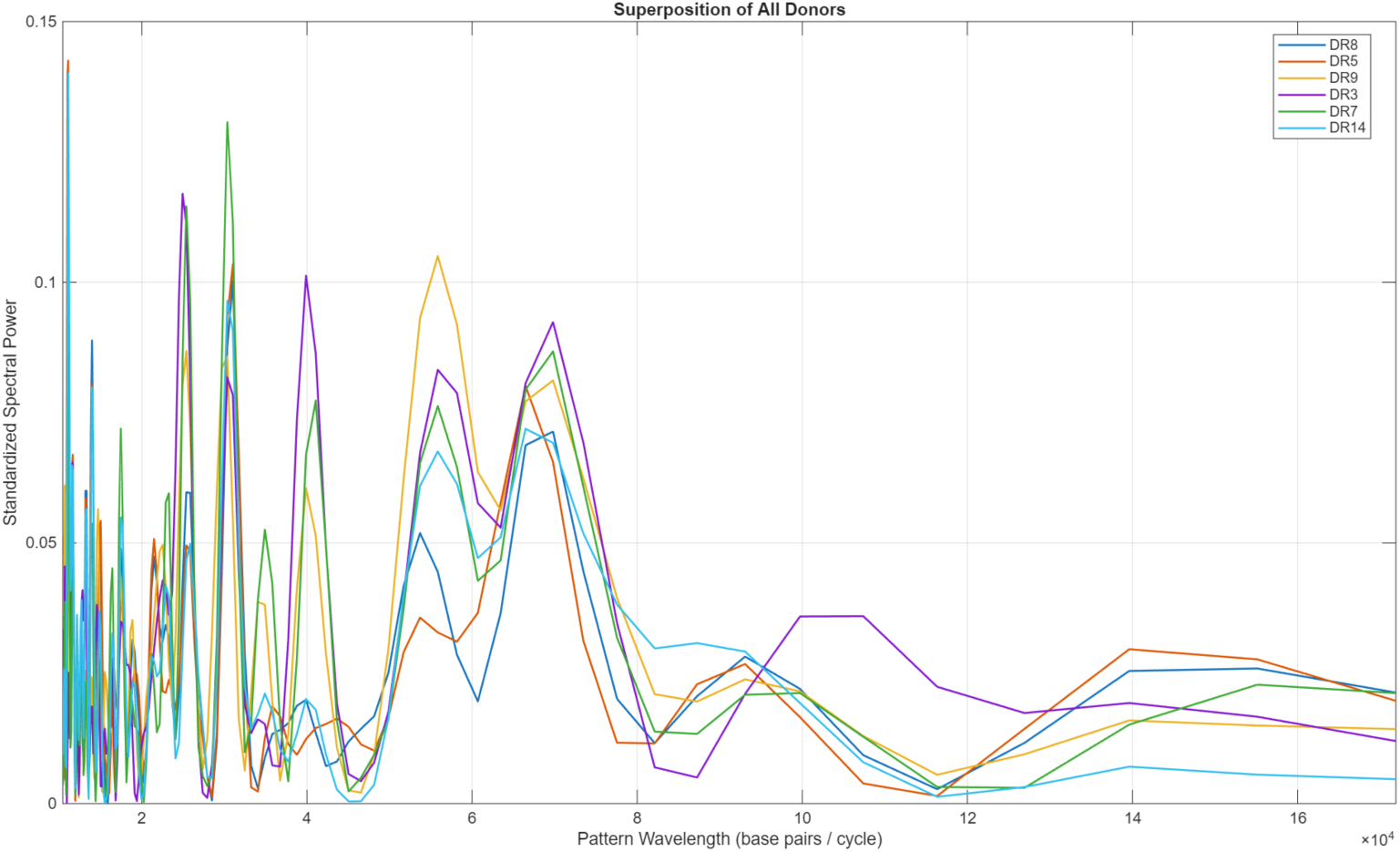
Donor Lomb-Scargle power spectra for all six donor-recipient pairs. The frequency axis spans 16.7 to 500 millicycles/kb. Power spectra are normalized to total power = 1. The 6 Donors show a conserved spectral signature with dominant power in the 110-250 millicycles/kb range (corresponding to 4-9 kb wavelength). Second panel shows that donor spectral wavelengths are consistent across normal donors in keeping with the frequency domain. Scale on the Y axis, is 10^4^ base pairs per cycle. The wavelength spectra depicted were calculated using MATLAB and are not subject to frequency binning utilized for the frequency spectra.

**Figure 2.**
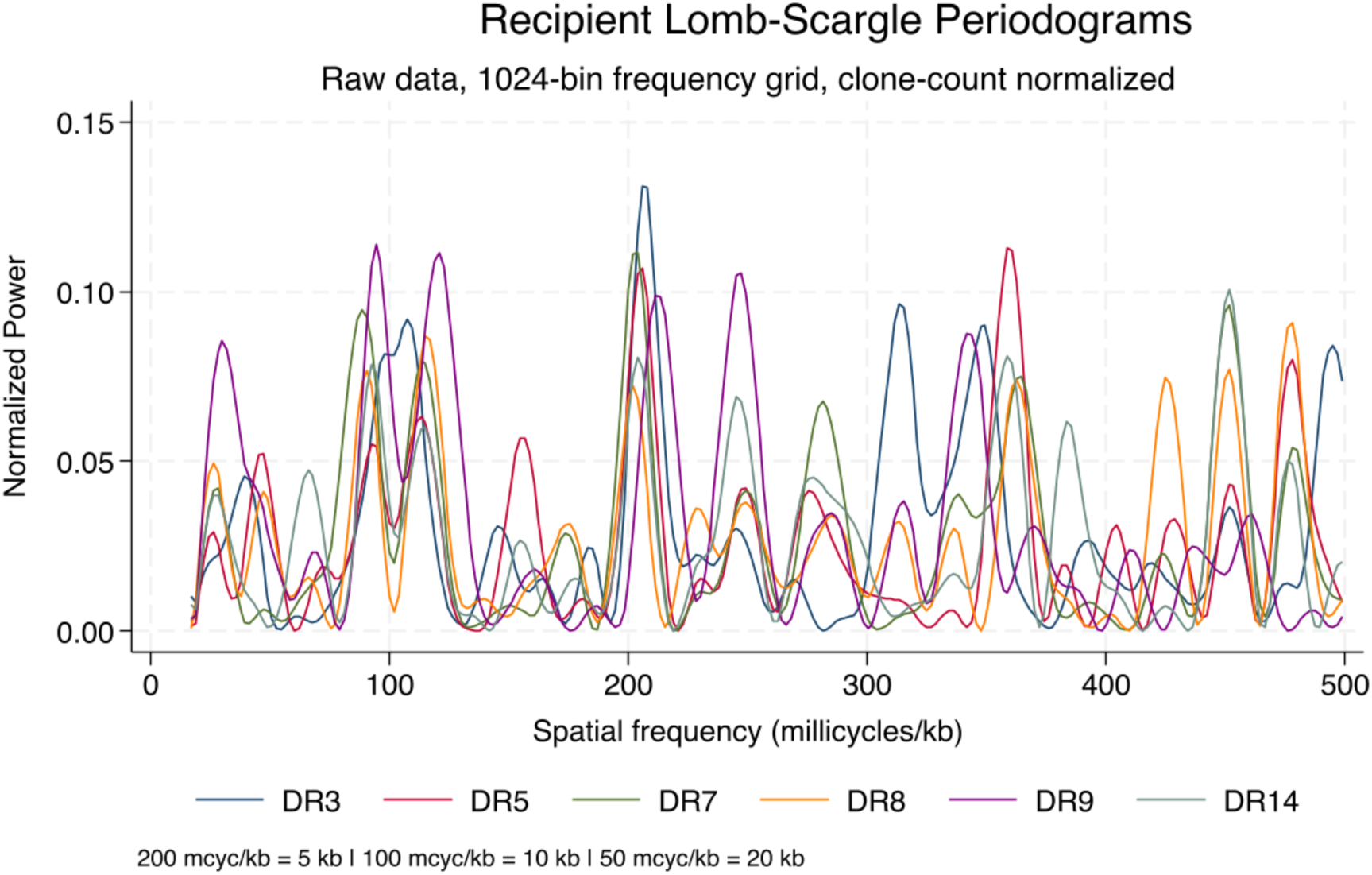
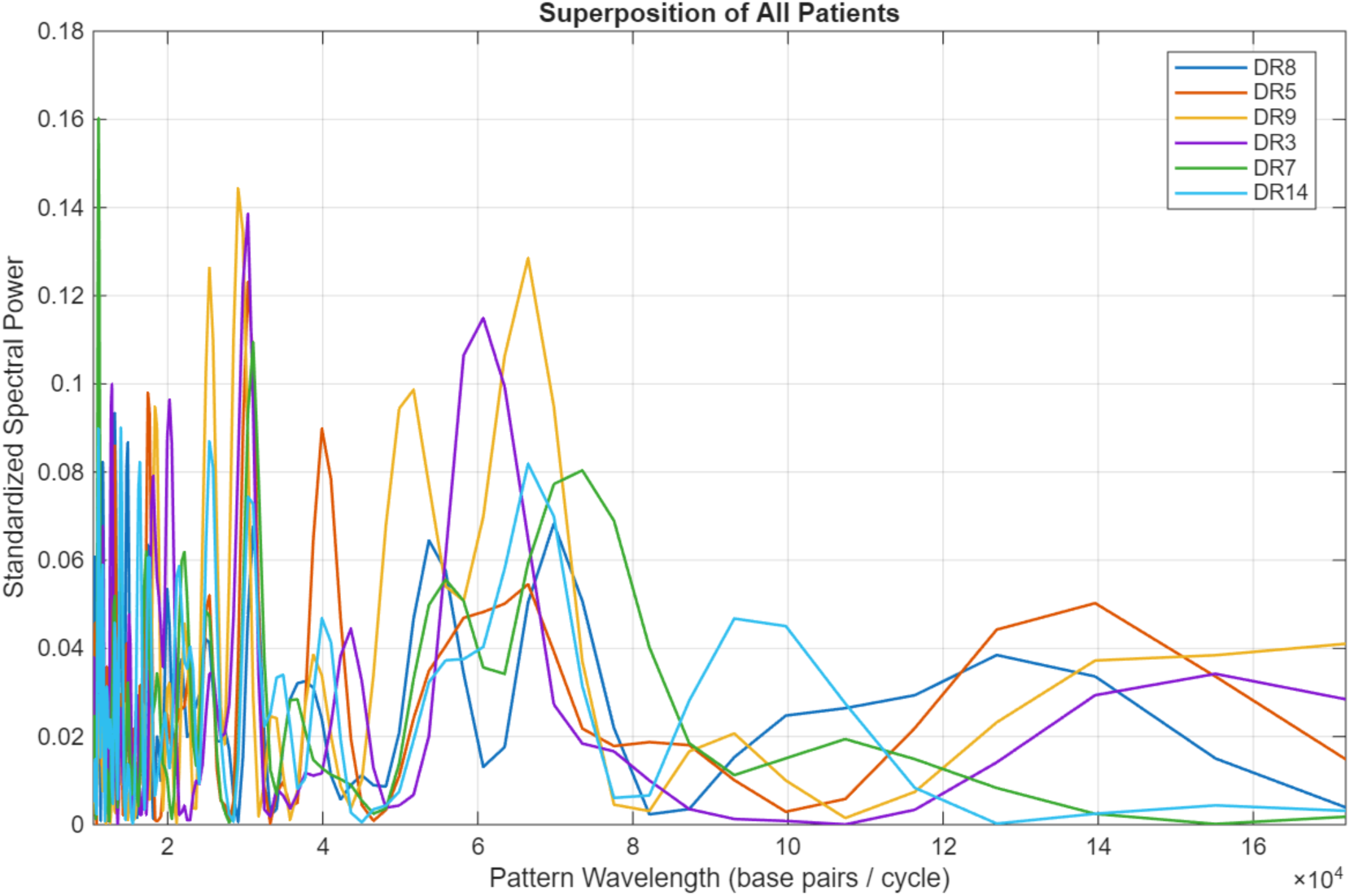
Overlay of all six recipient power spectra. While Donor spectra (Figure 1) show tight clustering, indicating a conserved spectral signature across individuals, Recipient spectra show greater dispersion, reflecting variable post-transplant reconstitution. Recipients diverge from their matched donors to varying degrees, shifting towards longer wavelengths.

### Primary Spectral Metrics

The primary spectral metrics computed for each sample are summarized in Table 1. The spectral centroid (SC) ranged from a relatively narrow range of 231.0 to 249.1 millicycles/kb in donors (median 240; ∼4 Kb wavelength), and a more dispersed, 213.6 to 264.9 millicycles/kb in recipients (median 260), reflecting greater disorder and randomness in the usage of the V gene segments in the post-transplant reconstituting T cell repertoire. In 5 of 6 pairs, the recipient’s SC was higher than the paired donor’s (**Table 1**, **Figure 3, Supplementary Table 1**), indicating a shift toward higher-frequency (shorter-wavelength) spectral components after transplantation. These findings support the notion, that the recipient T cell repertoire has disruption of the normal periodic distribution observed in the donor T cell repertoire V gene representation in the T cell clones and a more uniform representation of V segments in the repertoire. In other words, the normal hierarchy of V segment usage is disrupted in the post-transplant setting.

**Figure 3.**
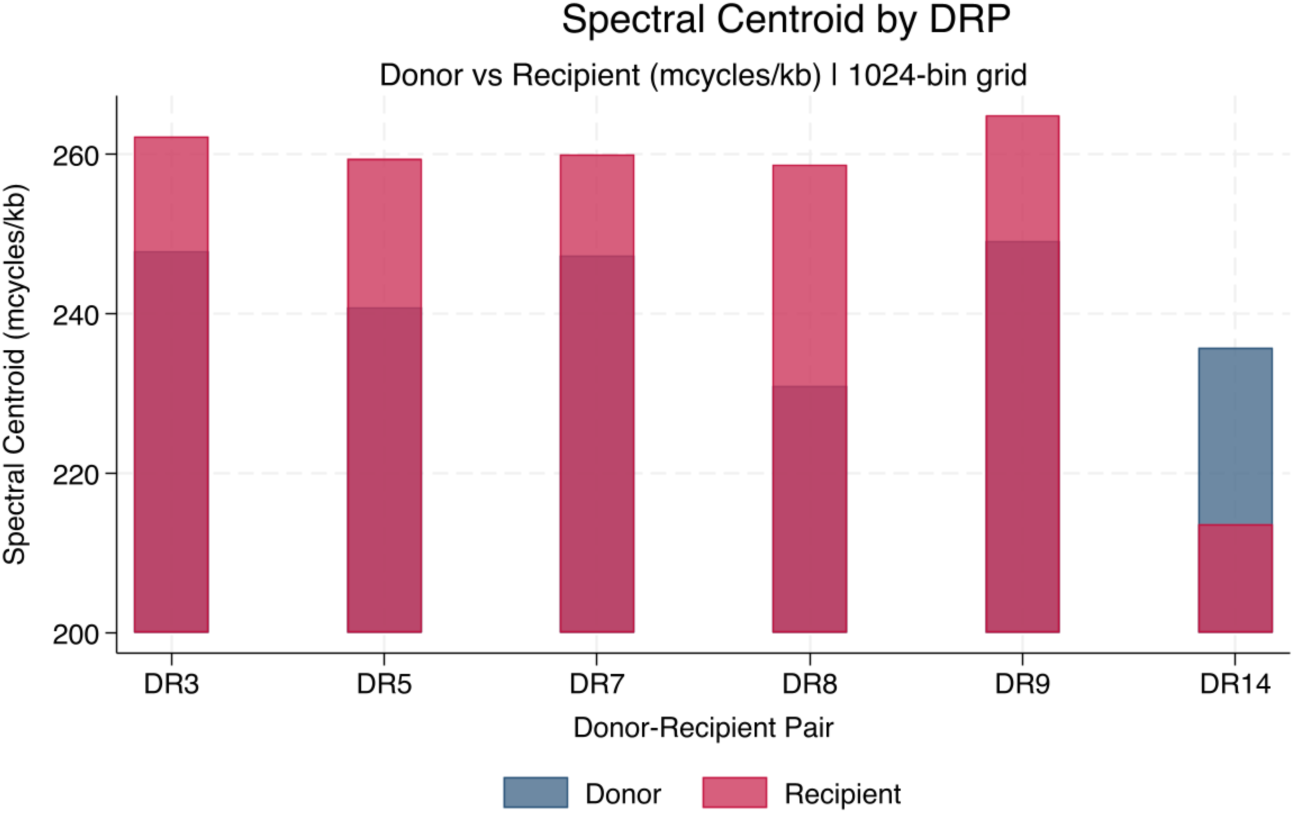
Spectral centroid comparison for each donor-recipient pair. Five of six pairs show a higher SC in recipients; band power is similar within pairs.

**Table 1.** Spectral Metrics per Donor-Recipient Pair. * DRP 9 with large decline in SC from D to R, DRP 3 and 7 with a large increase in R from D;

| Pair | SC_D | SC_R | Delta_SC | LFI_D | LFI_R | Ent_D | Ent_R |
| --- | --- | --- | --- | --- | --- | --- | --- |
| DR3 | 240.8 | 259.5 | +18.7 | 0.178 | 0.144 | 9.50 | 9.41 |
| DR5 | 247.3 | 260.0 | +12.7 | 0.224 | 0.163 | 9.50 | 9.43 |
| DR7 | 231.0* | 258.7* | +27.7 | 0.199 | 0.183 | 9.41 | 9.38 |
| DR8 | 249.1 | 264.9 | +15.7 | 0.215 | 0.175 | 9.45 | 9.53 |
| DR9 | 235.8 | 213.6 | -22.1 | 0.200 | 0.233 | 9.54 | 9.37 |
| DR14 | 247.8 | 262.3 | +14.4 | 0.215 | 0.180 | 9.43 | 9.47 |

*SC units: millicycles/kb. Delta = Recipient - Donor. LFI = low-frequency index (fraction of power <100 millicycles/kb). Ent = spectral entropy (bits). D/R\_raw = uncorrected donor/recipient total power ratio. D/R\_norm = clone-count normalized ratio (~1.0 if shape preserved).*

Spectral entropy ranged from 9.41 to 9.54 bits in donors and 9.37 to 9.53 bits in recipients, with no consistent donor-recipient direction (2 of 6 pairs showed increased entropy in recipients). The uniformly high entropy values (maximum possible for 1024 bins is approximately 10 bits) indicate that spectral power is broadly distributed across the frequency grid. This is likely a consequence of the normally complex structure of the T cell repertoire given the multiplicity of V genes involved under normal circumstances, and the afore mentioned complexity of the periodic V segment usage waveform.

### DRP Fourier Analysis

The recipient V gene expression patterns resembled the donors but with alterations in the frequency domain (**Figure 2**). Three of the recipients (DR8, DR7, and DR14) had spectra similar to their donors, with dominant wavelengths in the 5-11 kb range. Three others showed more pronounced divergence, particularly DR3, where the recipient spectrum showed the most extreme displacement. Recipients in DR8 and DR14 had higher entropies than their donors, reflecting a more disordered repertoire in the recipients in terms of relative TRB V gene segment usage. DR3 and DR9 recipients, on the other hand, demonstrated marked spectral changes consistent with diminishing diversity in the V gene segment utilization (**Table 1**). The dominant wavelengths observed in the recipients were similarly dispersed when compared with those in the donors (**Supplementary Figure 2**). As these analytic data are interpreted, it is important to recognize that these results are calculated and thus distinct from the measured variables they are derived from.

### Band Power Distribution

The most robust spectral finding is the distribution of power across biologically defined wavelength bands. Across all six pairs, the 3-12 kb band (sub-family spacing) accounts for 55.9 +/- 3.9% of total spectral power in donors and 54.9 +/- 4.8% in recipients (**Table 2 and Figure 4**) (**Supplementary Figure 2**). This confirms that the sub-family spacing pattern dominates the TRB V gene usage spectrum. The 12-30 kb cluster band accounts for 7.3 +/- 0.9% (donors) and 6.9 +/- 1.2% (recipients), while the 30-60 kb inter-cluster band contributes 4.9 +/- 1.2% and 3.6 +/- 1.4%, respectively. The residual high-frequency band (<3 kb) accounts for 32.0 +/- 2.4% (donors) and 34.5 +/- 5.8% (recipients). The mean donor-recipient difference in 3-12 kb band power was -1.0 percentage points (range: -6.2 to +4.5), with no consistent direction across pairs. The within-pair similarity of band power — despite individual differences in total clonal frequency and spectral centroid — suggests that the sub-family spacing pattern is a conserved property of the TRB repertoire architecture rather than a marker of repertoire perturbation.

**Figure 4.**
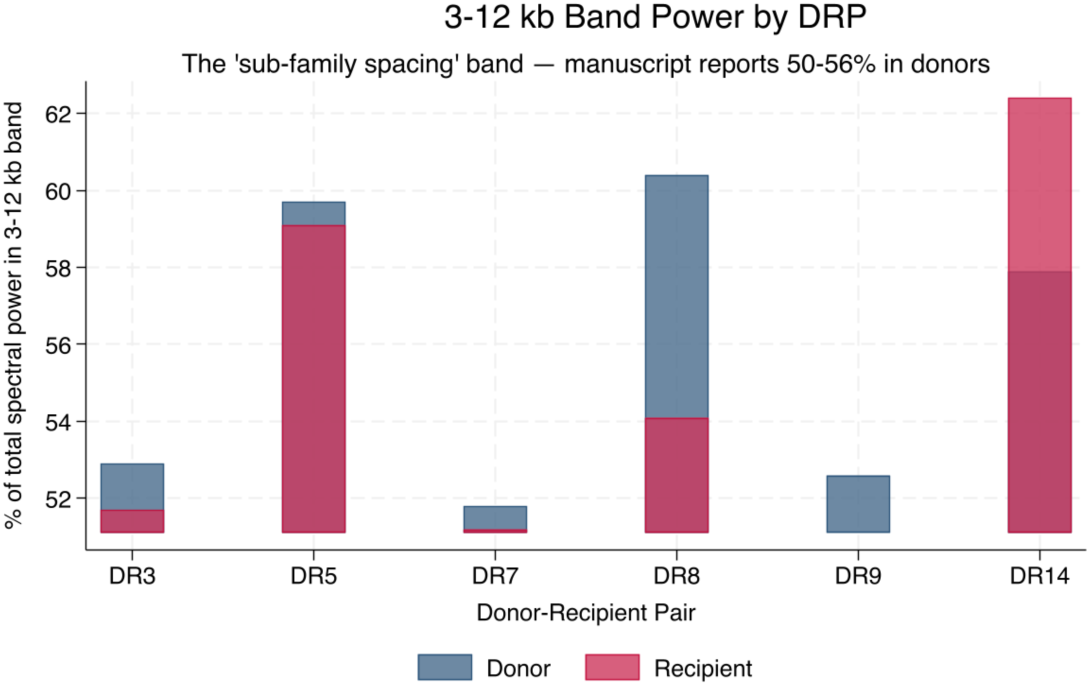
3-12 kb band power comparison between donors and recipients.

**Table 2.**
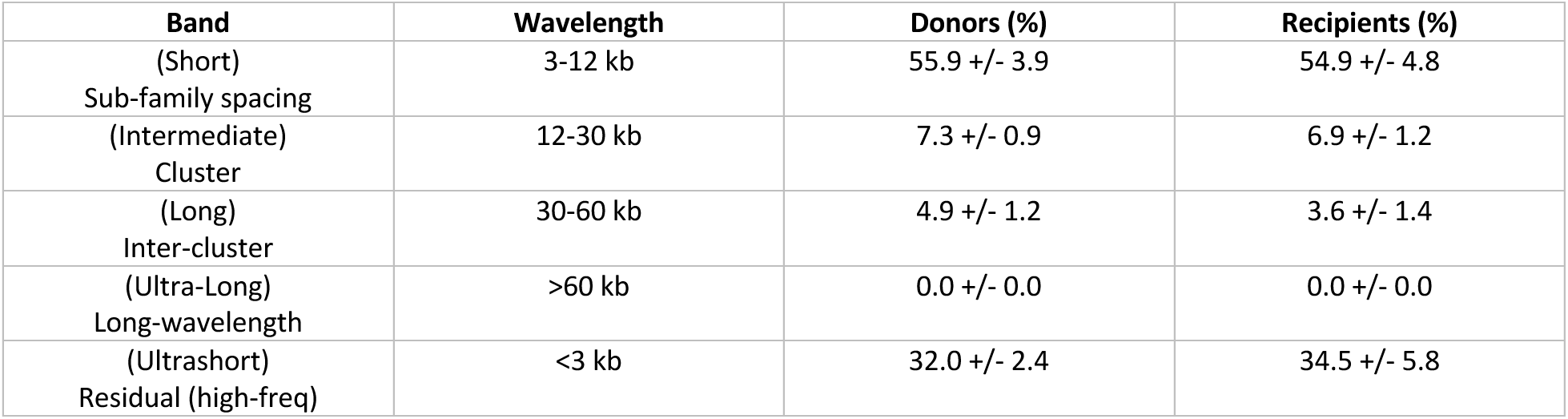
Spectral Power Distribution by Wavelength Band (Mean +/- SD, n = 6 Pairs)

| Band | Wavelength | Donors (%) | Recipients (%) |
| --- | --- | --- | --- |
| (Short)<br>Sub-family spacing | 3-12 kb | 55.9 +/- 3.9 | 54.9 +/- 4.8 |
| (Intermediate)<br>Cluster | 12-30 kb | 7.3 +/- 0.9 | 6.9 +/- 1.2 |
| (Long)<br>Inter-cluster | 30-60 kb | 4.9 +/- 1.2 | 3.6 +/- 1.4 |
| (Ultra-Long)<br>Long-wavelength | >60 kb | 0.0 +/- 0.0 | 0.0 +/- 0.0 |
| (Ultrashort)<br>Residual (high-freq) | <3 kb | 32.0 +/- 2.4 | 34.5 +/- 5.8 |

### Donor-Recipient Comparison

Every donor in every dataset shows its dominant spectral power concentrated at spatial wavelengths of 4-9 kb, corresponding to spatial frequencies of approximately 113-252 millicycles/kb. Recipients diverged from this markedly across the frequency spectrum (**Figure 5**), implying differential V segment usage in the recipient as the T cell repertoire reconstitutes following transplantation. Recipients diverged from this in a direction that depends on their recovery state. Collapsed and contracted recipients (DR3, DR9) showed more pronounced spectral alterations, while some recipients (DR7, DR14, DR8, DR5) retained dominant wavelengths in the 5-11 kb range but with altered power distribution.

**Figure 5.**
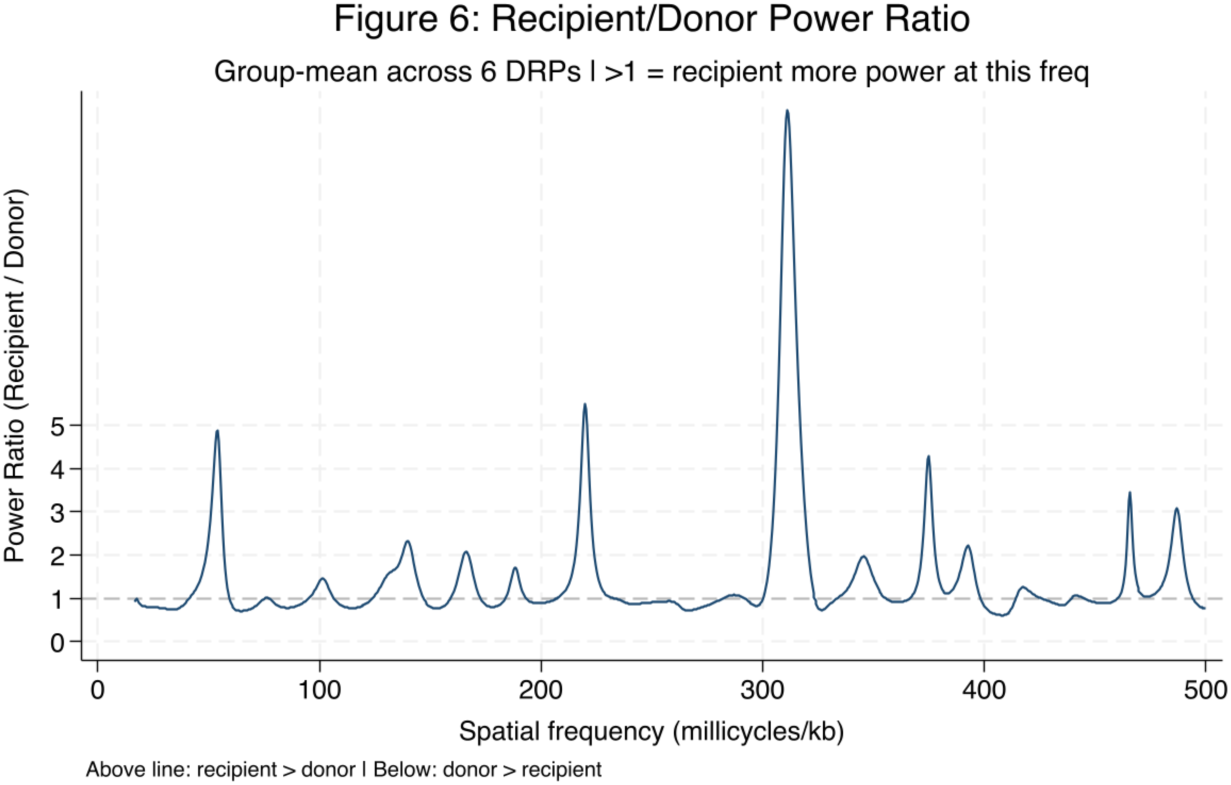
Donor-recipient power ratio across the frequency spectrum. Values >1 indicate relatively more power in the recipient at that frequency. Recipients carry approximately 2-3x more power than donors at very short wavelengths (3-5 kb) and at 20-40 kb.

Consistent with the higher frequency shift in the spectral centroid, the low frequency index was lower in the recipients as well (**Figure 6**).

**Figure 6.**
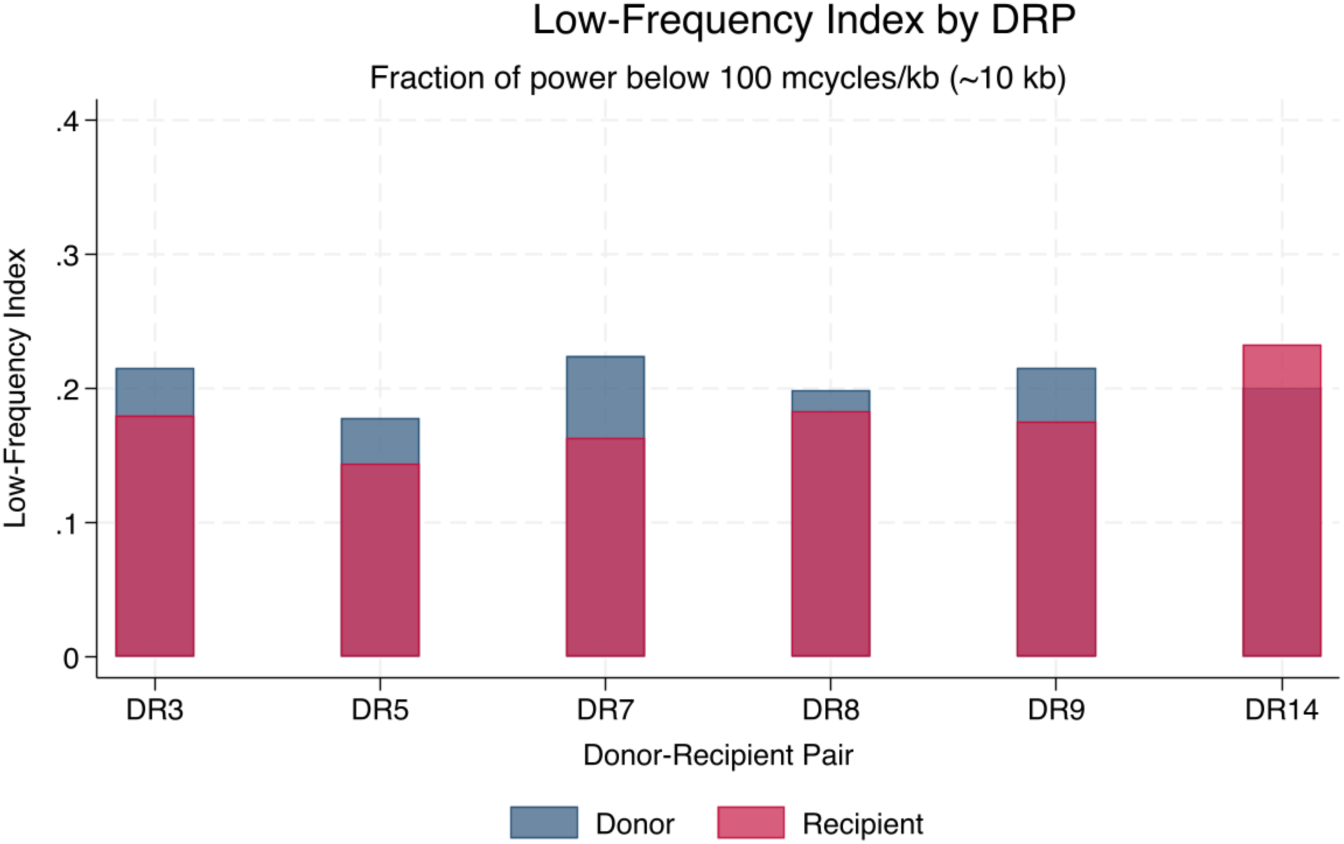
Low-frequency index comparison and band power comparison highlighting the distribution of spectral energy across wavelength bands.

### Sensitivity Analysis

The 3-12 kb band power was computed across three grid sizes (128, 512, and 1024 bins) at two frequency cutoffs (f_min_ = 1/span and f_min_ = 1/60 kb) (Table 4). At a fixed f_min_, increasing the number of bins from 128 to 1024 changed the band power by less than 2 percentage points in all pairs, demonstrating that the band-level metric is essentially invariant to grid resolution once a reasonable minimum is reached.

**Table 4.** Sensitivity Analysis — 3-12 kb Band Power in Donors Across Six Settings

| Setting | DR3 | DR5 | DR7 | DR8 | DR9 | DR14 |
| --- | --- | --- | --- | --- | --- | --- |
| 128 bins,<br>$f_{\min}=1/\text{span}$ | 48.3% | 42.7% | 52.7% | 43.5% | 48.4% | 44.9% |
| 512 bins,<br>$f_{\min}=1/\text{span}$ | 49.6% | 44.0% | 53.9% | 44.7% | 49.6% | 46.0% |
| 1024 bins,<br>$f_{\min}=1/\text{span}$ | 49.9% | 44.2% | 54.1% | 45.0% | 49.8% | 46.3% |
| 1024 bins,<br>$f_{\min}=1/120$ | 55.2% | 48.8% | 58.1% | 49.4% | 54.4% | 50.4% |
| 1024 bins,<br>$f_{\min}=1/60$ * | 59.7% | 51.8% | 60.4% | 52.6% | 57.9% | 52.9% |
| 128 bins,<br>$f_{\min}=1/60$ | 59.7% | 51.9% | 60.6% | 52.7% | 58.1% | 53.1% |

\* Primary analysis setting. The near-identity of the last two rows (mean delta = 0.15 pp) confirms that the standardized $f_{\min} = 1/60$ kb, rather than grid resolution, is the primary determinant of band power.

Raising f_min_ from 1/span (∼0.0029 cycles/kb, lambda _max_ ∼344 kb (wavelength)) to 1/60 kb (0.0167 cycles/kb, lambda _max_ = 60 kb) increased the 3-12 kb band power by 6-10 percentage points across pairs. This shift is expected: excluding the very low frequencies (wavelengths >60 kb) reallocates that power to the remaining bands, primarily the 3-12 kb and <3 kb bands. The manuscript’s reported 3-12 kb band power of 50- 56% is reproduced at the standardized f_min_ = 1/60 kb (51.8-60.4% in our analysis), confirming that the earlier discordance with the manuscript was entirely a function of the frequency window, not a biological difference.

### Jensen-Shannon Divergence

The Jensen-Shannon Divergence, which quantifies the overall dissimilarity of the normalized power spectra between paired donors and recipients, ranged from 0.18 (DR14) to 0.35 (DR3) in natural units (**Table 5**). The ranking of pairs by JSD is: DR3 > DR9 > DR5 ∼ DR7 > DR8 > DR14. This ranking suggests that JSD captures variation in repertoire perturbation even when individual spectral metrics lack statistical significance. The Kullback-Leibler (KL) divergences were roughly symmetric in all pairs (KL(D to R) ∼ KL(R to D)), indicating that the spectral differences are not systematically directional in one condition. This is consistent with the similar band power distributions between donors and recipients.

**Table 5.** Jensen-Shannon Divergence between paired spectra.

| Pair | JSD (natural) | JSD (log2) |
| --- | --- | --- |
| DR3 | 0.3461 | 0.4157 |
| DR5 | 0.2609 | 0.3133 |
| DR7 | 0.2378 | 0.2856 |
| DR8 | 0.2120 | 0.2546 |
| DR9 | 0.2849 | 0.3421 |
| DR14 | 0.1839 | 0.2208 |

*JSD = 0 indicates identical spectra; higher values indicate greater divergence.*

## Discussion

In patients with hematological malignancies, an allogeneic SCT is inherently dependent on the donor-recipient alloreactivity for its therapeutic graft vs. leukemia effect, however often this is complicated by graft vs. host disease. Immune suppression impedes T cell recovery, which in turn increases the risk for infection, as does the oligoclonal expansion of alloreactive T cells post-transplant. These effects impact donor-recipient pairs stochastically and are not predictable with a high degree of accuracy, given the heterogeneity inherent to the HLA system, and the human exome as a source of alloreactive minor histocompatibility antigens.^7^ T cells are the main protagonists for these immunological phenomenon and deep analysis of the T cell repertoire with NGS has been utilized to help understand immune recovery post-transplant and its influence on clinical outcomes.

However, the great variability observed in the T cell repertoire which is generated by VDJ recombination and untemplated nucleotide addition to the CDR3 region of TRA and TRB makes it difficult to correlate changes in the repertoire with clinical outcomes. Numerous summary measures such as Shannon’s entropy, Euclidean distance and T cell clone tracking strategies have been developed to quantify the differences between donor and recipient T cell profiles, but none have thus far yielded a consistently reliable measure of alloreactivity/immune failure, often because the effects are mediated by polyclonal repertoire shifts occurring in the broader context of numeric T cell population changes. These polyclonal cross repertoire changes are not necessarily fully captured by any of the summary methods currently utilized.

In this paper, a technique widely utilized in signal processing is applied to understand T cell repertoire changes following transplantation, using the donor to recipient transitions in T cell repertoire as a model system. Fourier analysis transforms periodic data in time or spatial domains to the frequency domain, deconstructing complex signals into the constituent frequencies to enable identification of the most important signal components which carry the greatest information or Power in a signal. In essence, transforming the original periodic data into frequencies reveals which wavelength/frequency carries the most information versus trivial details or noise. As an example, in day-to-day life, this helps compress information and in data storage. In evaluating the T cell repertoire, the relative frequencies of the T cell receptor beta (or alpha) V or J segment defined clones may be similarly considered. These TRB and TRA, VDJ recombined sequences, are generated by genomic loci which are logarithmically scaled and periodically distributed. Previous work has demonstrated that spatial ordering the T cell clonal frequencies according to the location of the V or J segments loci on the TRB locus reveals a periodic pattern which is well organized and near identical in pattern in all SCT donors studied, but this ordering is disrupted in SCT recipients. Visually the magnitude of disruption is greater in those afflicted by GVHD, or infections due to poor immune recovery, but no reliable, objective measure has emerged to help quantify the magnitude of disruption in the T cell repertoire from donor to recipient, across V or J gene segment defined clones, or assess the status of an individual’s immune competence compared with a population normal.

Fourier analysis of T cell receptor beta V gene defined clonal frequencies examined as a function of the location of the relevant gene segment on the TRB locus reported herein, demonstrates that within the frequency domain, the normal stem cell donors had a consistent and reproducible spectral pattern of V gene segment usage, even though these were unrelated individuals. This implies that there is broadly similar use of the TRB V gene segments to yield a diverse, polyclonal repertoire in normal individuals. Furthermore, the relative representation of various V gene segments in the repertoire is dependent on the location of the V (and J) gene segments on the loci. The longer wavelength of spectral centroid and greater value of low frequency indices (LFI) in the donors suggests an organized TCR repertoire, with a hierarchy of V gene segments involved in the recombination process and clonal generation. This organization has been previously demonstrated in the observations on the Fractal nature of the T cell repertoire. On the other hand, recipients had a more varied and diffuse power distribution across a broader range of spatial frequencies. The spectral centroid had a higher frequency in general and correspondingly, the LFI was lower. This is consistent with a more chaotic and disorderly utilization of TRB V gene segments in the recipient, a less hierarchical VDJ recombination. These polyclonal changes may be attributable to different T cell clones expanding disproportionately and others failing to recover adequately under conditions of alloreactive and pathogen pressure, as well as immune suppression. Eventual T cell reconstitution and in vivo TCR recombination will in such situations be expected to be restorative of the Power spectra to normal. In summary, one may posit that the donor frequency spectra are a conserved genomic baseline — a consistent fingerprint of the TRB locus physical structure under conditions of polyclonal normal recombination — while recipient frequency spectra are perturbations of that baseline whose direction and magnitude reflect the clinical state of T-cell reconstitution. The direction of the dominant wavelength change (stable, consistent with preserved repertoire breadth, versus, elongation indicating sub-family collapse); similarly in larger cohorts shifts in entropy and the band power redistribution from intermediate to short and long wavelengths together constitute a spectral fingerprint of immune recovery or response that is quantifiable, and not detectable by absolute clone counts or conventional diversity metrics alone.

The major advantage of this analytic methodology is that it provides a way to objectively quantify the robustness of the repertoire recovery, by potentially focusing on the usage of all the TRB gene segments instead of reporting on individual clones, as most clone tracking methods would. It also preserves more information about the T cell repertoire than a single measure such as Shannon’s entropy or Euclidean distance. Utilizing 2- dimensional Fourier Transforms, both the V and J gene segments may be analyzed. But most importantly, the consistency across the frequency spectra supports the dynamical systems nature of T cell responses, which occur in proportion to antigen exposure across the continuum of T cell repertoire.

Given the small sample size analyzed here and the relatively late time of sample acquisition, no conclusions may be drawn regarding the clinical utility of this analytic technique, but given the uniformity observed in the donors, and the dispersion seen in the recipients post-transplant, it is likely that patient with GVHD or CMV reactivation characterized by oligoclonal T cell proliferation will have disparate spectral signatures and when measured in the right context this will be a useful tool for discriminating these alloreactive states.

Thus, by changing the final pattern of the TRB V gene frequencies, relative expansion of certain clones in the recipient will alter the Power distribution across the constituent frequencies. This may allow discrimination of different clinic states between patients. Furthermore, this will likely be informative in other clinical situations where immune deficiency or oligoclonal autoreactive T cell expansion is a concern.

This is an initial proof of concept work and there is much room to improve upon the analytic pipeline presented here, in terms of the best frequency binning for the FFT (recognizing that ideally it is an infinite continuum), optimization of downstream analytics such as Spectral Entropy, Low Frequency Indices, appropriate scaling of the spatial parameters (whether Kb or Radians), incorporation of the J segment information. A larger cohort with sequencing based on contemporary TRA and TRB clonal definition will be best to accomplish this.

Additionally, further work on elucidating the biologic, structural basis of the dominant wavelengths observed in the Power Spectra will be of great interest in terms of the underlying biologic principles governing the evolution of immune responses.

## Acknowledgements

Artificial intelligence was used for the initial data analysis and proof of concept. Financial disclosures – The authors declare no relevant financial disclosures.

## Author attribution

AT: Developed the concept, organized and analyzed the data, wrote the paper; AMV: Analyzed the data and wrote the paper; RQ: Analyzed the data and wrote the paper.

## Supplementary Results

### Pseudogene Comb-Filter Test

Eighteen TRB V genes in this dataset are pseudogenes (marked with asterisks), having clonal frequency = 0 but occupying biologically relevant genomic positions. To test whether these zero-frequency data points introduce a periodic artifact (comb filter) into the power spectrum, the full pipeline was re-run after excluding all 18 pseudogenes, retaining 49 functional V genes.

Excluding pseudogenes changed the 3-12 kb band power by +0.6 to +1.1 percentage points across pairs (**Supplementary Table A**) and the spectral centroid by less than 0.7 millicycles/kb in all cases. The spectral fingerprint is therefore not driven by pseudogene artifacts and reflects genuine V-gene usage patterns in functional genes.

**Supplementary Table A.** Pseudogene Exclusion — Effect on 3-12 kb Band Power

| Pair | All 67 genes (%) | No pseudogenes (49 genes, %) | Delta (pp) |
| --- | --- | --- | --- |
| DR3 | 59.7 | 60.3 | +0.6 |
| DR5 | 51.8 | 52.8 | +1.0 |
| DR7 | 60.4 | 61.3 | +1.0 |
| DR8 | 52.6 | 53.7 | +1.1 |
| DR9 | 57.9 | 58.6 | +0.7 |
| DR14 | 52.9 | 53.9 | +1.0 |

### V30 Inclusion/Exclusion Sensitivity

Our primary analysis includes V30 for completeness. To verify that this does not affect the results, the pipeline was re-run excluding V30 (retaining 66 V genes). Excluding V30 changed the 3-12 kb band power by -0.2 to -0.9 percentage points, and the SC by -1.5 to -4.3 millicycles/kband JSD by -0.010 to +0.011 (**Supplementary Table B**). None of these changes are analytically meaningful, and the JSD ranking of pairs (DR3 > DR9 > DR5 > DR7 > DR8 > DR14) is preserved.

**Supplementary Table B.** V30 Exclusion — Effect on Key Spectral Metrics

| Pair | BP all (%) | BP no V30 (%) | Delta BP | SC all | SC no V30 | JSD all | JSD no V30 |
| --- | --- | --- | --- | --- | --- | --- | --- |
| DR3 | 59.7 | 59.5 | -0.2 | 240.8 | 238.6 | 0.3461 | 0.3572 |
| DR5 | 51.8 | 51.2 | -0.6 | 247.3 | 244.2 | 0.2609 | 0.2666 |
| DR7 | 60.4 | 60.1 | -0.3 | 231.0 | 226.7 | 0.2378 | 0.2279 |
| DR8 | 52.6 | 52.2 | -0.4 | 249.1 | 246.8 | 0.2120 | 0.2177 |
| DR9 | 57.9 | 57.6 | -0.3 | 235.8 | 234.3 | 0.2849 | 0.2844 |
| DR14 | 52.9 | 52.0 | -0.9 | 247.8 | 244.2 | 0.1839 | 0.1885 |

**Supplementary Figure 1.**
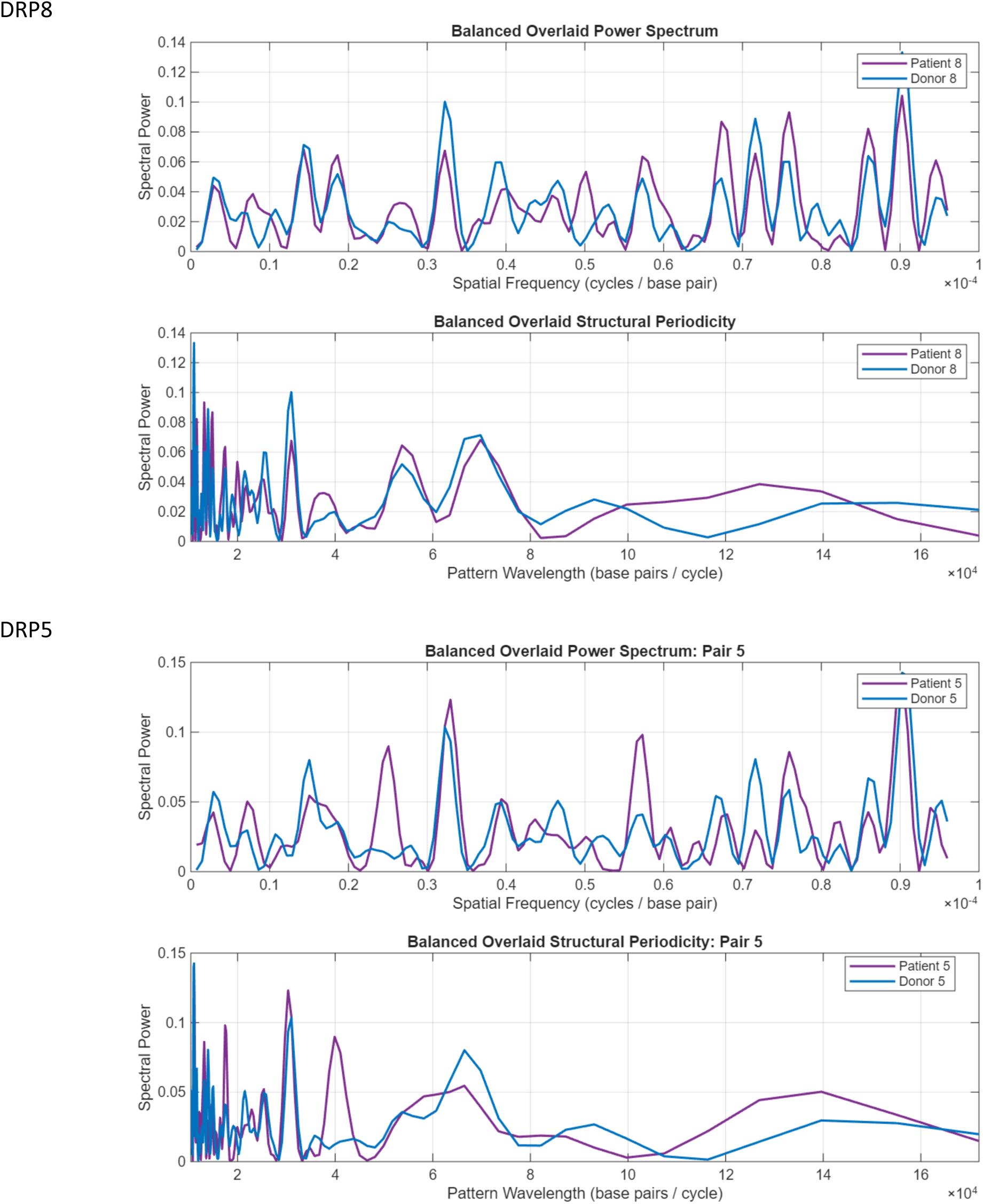
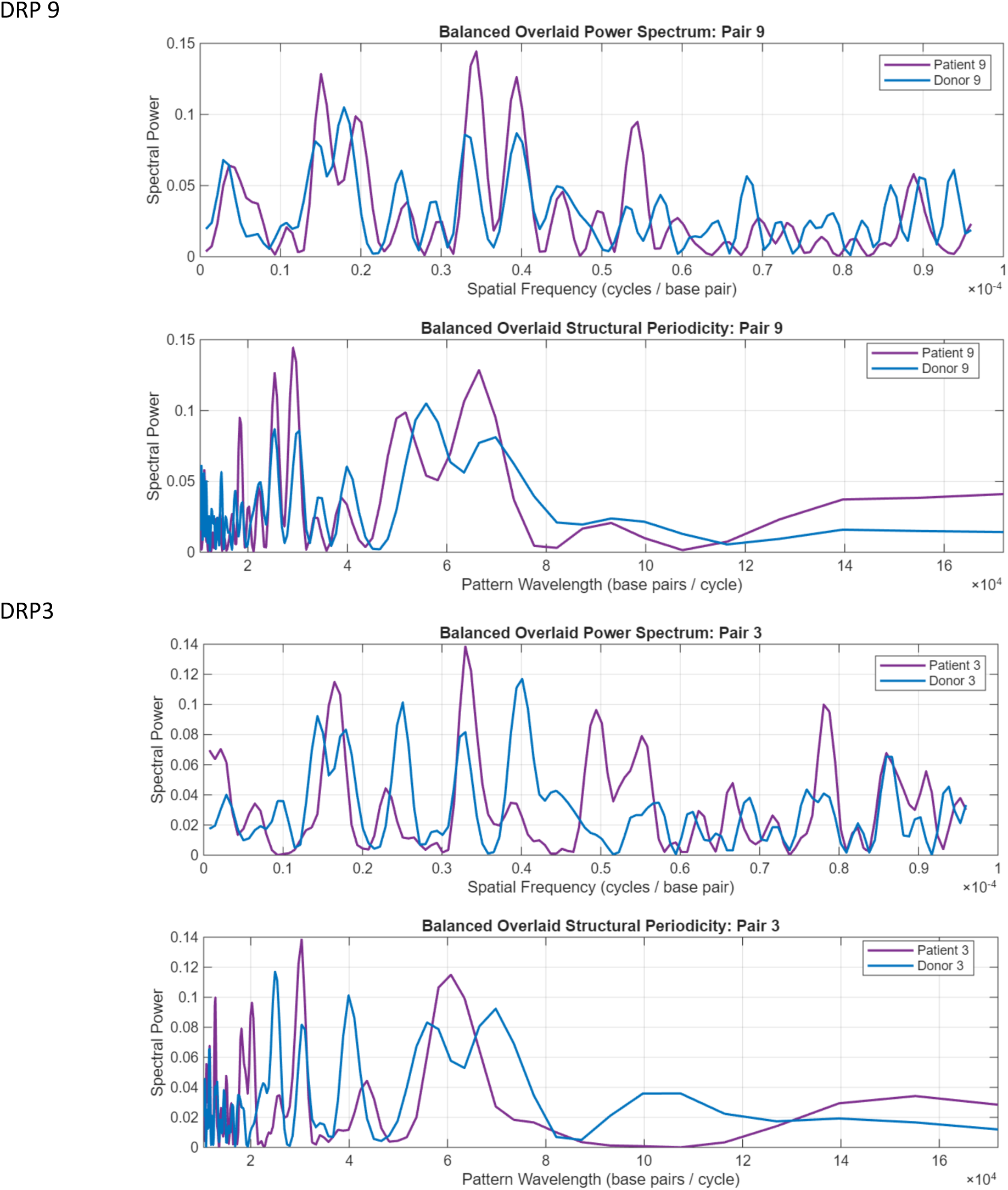
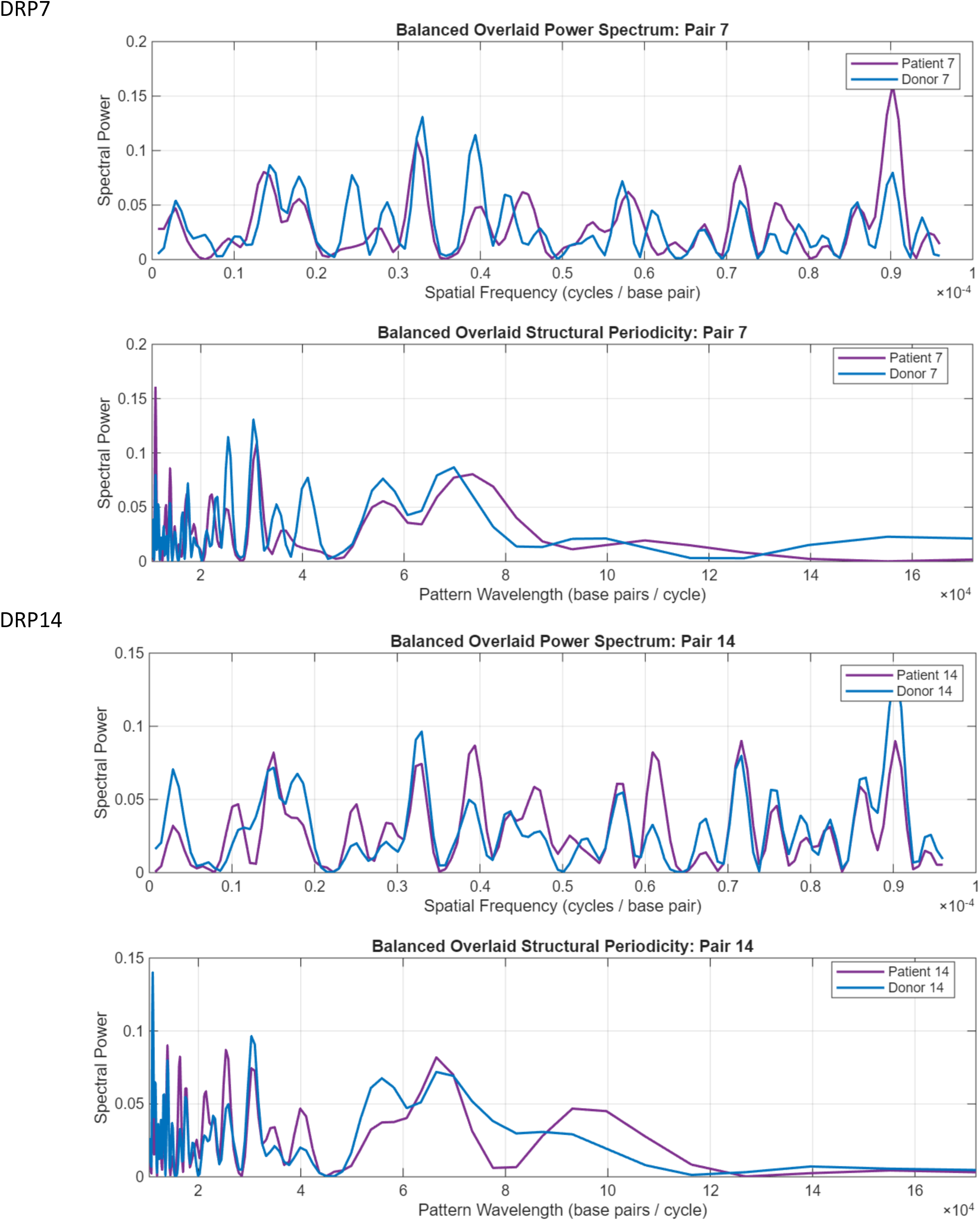
Comparison of individual DRP Power spectra; Spatial frequency 0.1*10^-4^ cycles/bp = 0.1*10^-1^ cycles/kb. (MATLAB spectral analysis output)

**Supplementary Figure 2:**
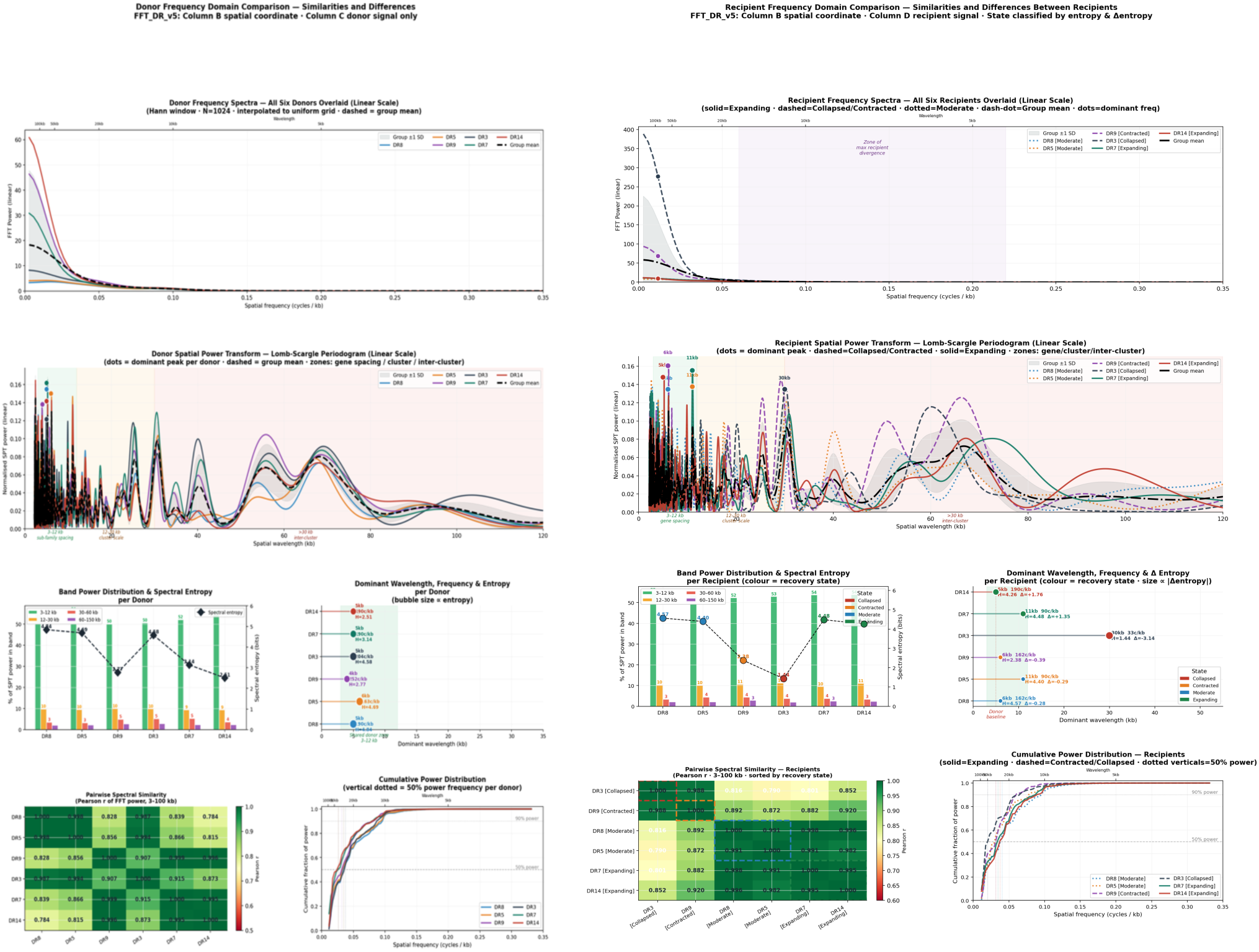
Comparison of donor and recipient characteristics generated by Claude.ai.

**Supplementary Table C.** Spectral parameters calculated using MATLAB without frequency binning. Spectral centroid and entropy behaves similarly as in frequency constrained calculations, with a shift towards higher frequencies and larger range of values observed in the recipients.

|  | Rec. Centroid | Don. Centroid | Centroid dif | Rec_ENT | Don_ENT |
| --- | --- | --- | --- | --- | --- |
| DR8 | 0.000053 | 0.000052 | -0.000001 | 6.6255 | 6.6351 |
| DR5 | 0.000051 | 0.000053 | 0.000002 | 6.5165 | 6.6307 |
| DR9* | 0.000038 | 0.000043 | 0.000005* | 6.3584 | 6.6867 |
| DR3* | 0.000048 | 0.000044 | -0.000004* | 6.4674 | 6.6053 |
| DR7* | 0.000051 | 0.000044 | -0.000007* | 6.5319 | 6.5462 |
| DR14 | 0.000050 | 0.000051 | 0.000001 | 6.5708 | 6.5799 |

## Appendix

The AI resources were queried to perform Fourier analyses and Fast Fourier Transform calculation on TCR clonal frequency data sets to quantify the variation between relative V gene usage in the donors and recipients. The distribution of TRB V defined clonal frequency was periodic in the spatial domain (Supplemental Figure 1) (Claude.ai). FFT analysis of the spatial domain, periodic graph of V gene copy numbers, decomposed these data into constituent frequency spectra (frequency domain), measuring the contribution of each frequency to the final configuration of the periodic curve in the spatial domain, a measure termed Spectral Power. The frequency spectra for donors and recipients were considered separately and then compared across the two groups.

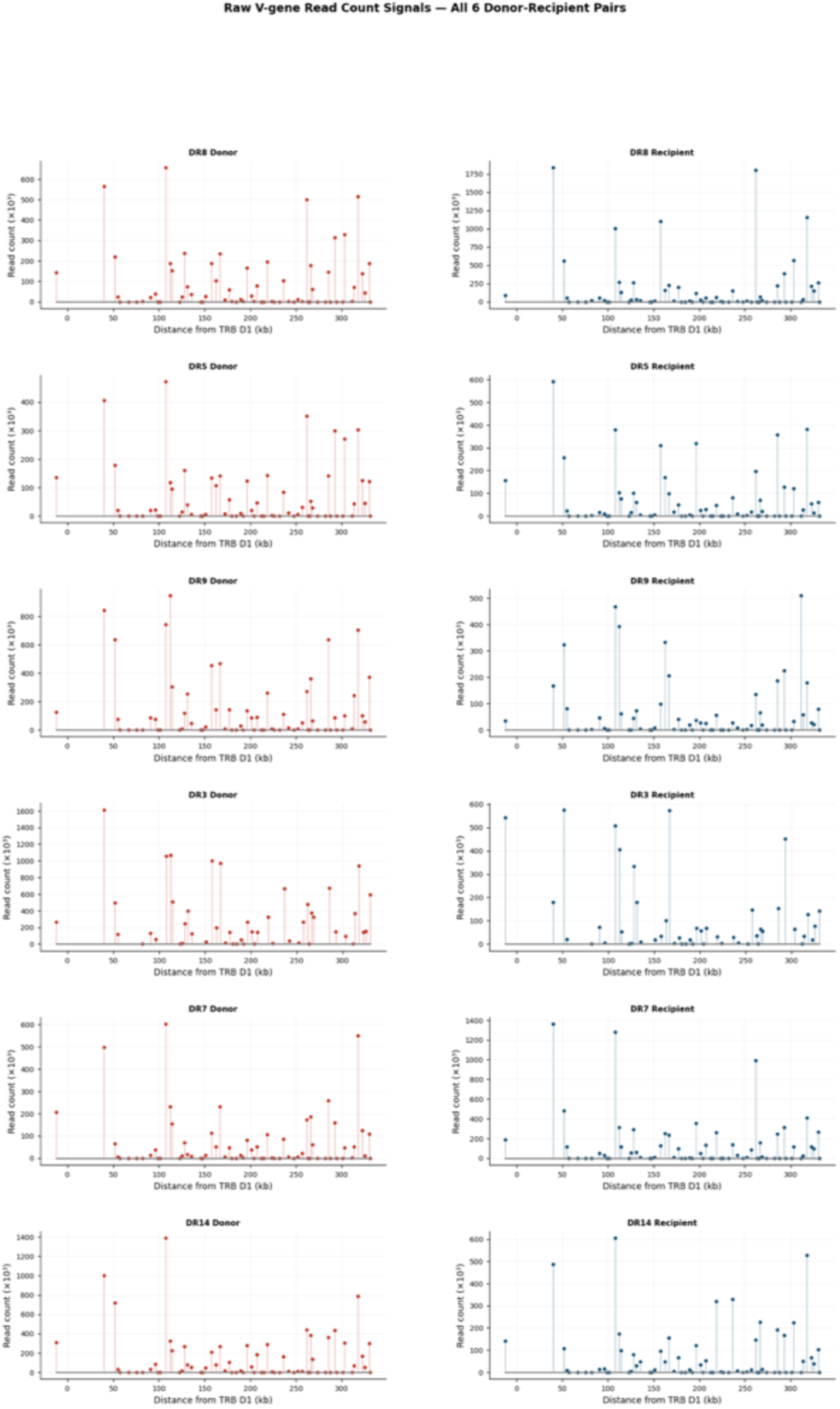

### Donor Fourier Analysis

The primary data analyzed were the copy number of TRB V segment defined T cell receptors mapped to the V gene segments on the TRB locus, in other words magnitude (amplitude) plotted in a spatial domain. As previously reported, an oscillating periodic output was observed consistently across all donor samples. While there was some variation observed in the magnitude of the oscillating wave, the pattern of relative V gene segment incorporation in TCR was similar across the donors. Fourier analysis in Claude demonstrated that there were common dominant spatial frequency bands, corresponding to a wavelength between 3-12 Radians (∼KB).

All six donors show 50–56% of their Lomb-Scargle power concentrated in this sub-family spacing band, and all have dominant wavelengths tightly clustered at 4–6 kb (163–252 cycles/kb) (Figure 1, Panel A, B and D). This may be considered a healthy-donor spectral signature. The group mean spectrum (dashed black, Panels A and B) shows a peak at ∼5 kb with a rapid decay into higher wavelengths.

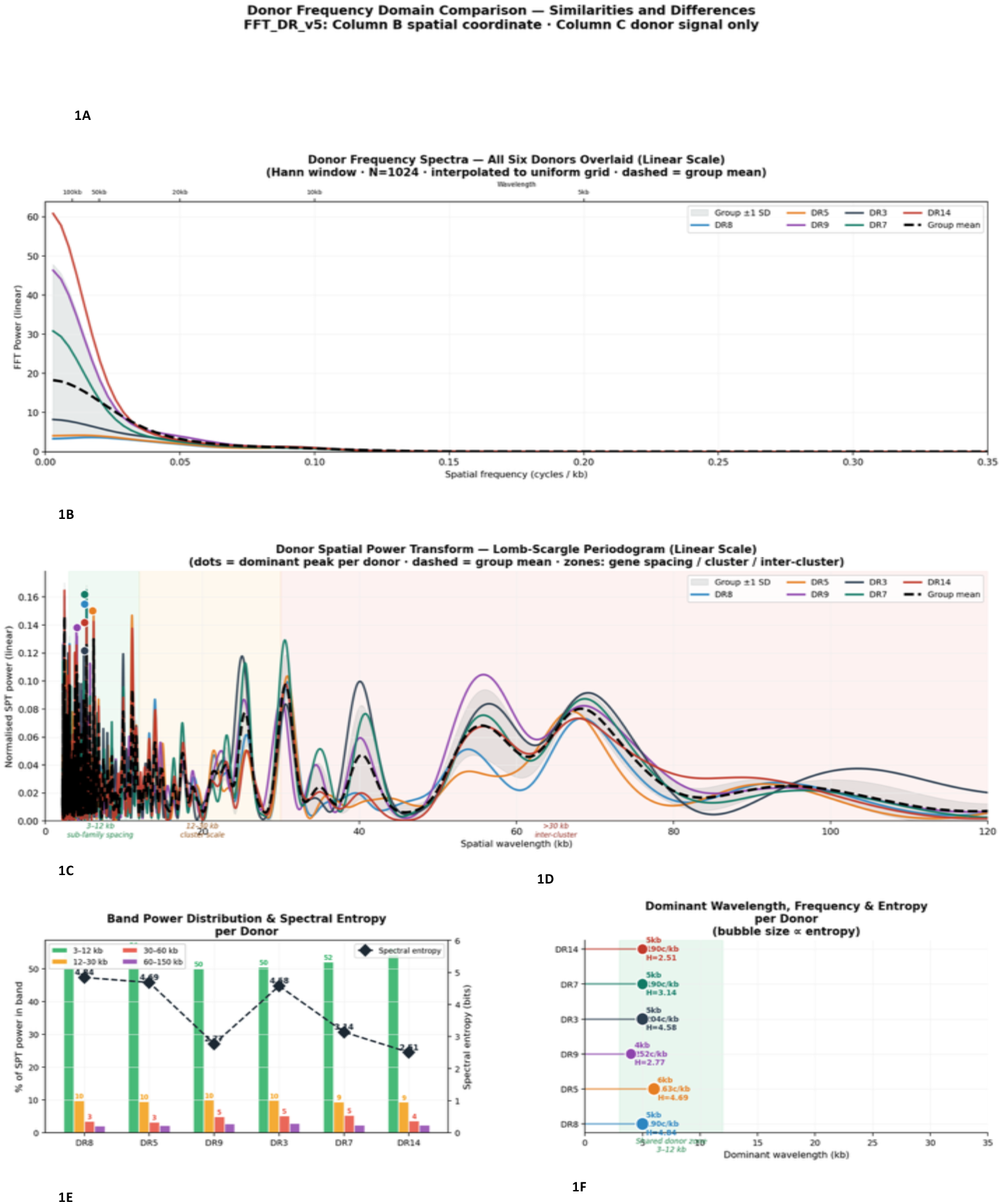

### Recipient Fourier Analysis

Next the recipient repertoires were analyzed to determine the characteristics of their frequency domain spectra. The recipient V gene expression patterns resembled the donors but with a shift towards longer wavelengths in the frequency domain as opposed to the relatively preserved V gene distribution of the donors, such that where donors clustered into fairly similar subgroups, the recipients spanned a range of spectral states (Figure 2A and 2B). Three of the recipients DR 8, 7 and 14 had similar spectra, with dominant wavelength of 5-11 radians (∼kB), three others had longer wavelengths, particularly the recipient in DR3.

The dominant wavelengths had a shift towards longer end of the spectrum consistent with loss of T cell clones utilizing successive V gene segments in a more uniform manner than the normal donors (Figure 2D).

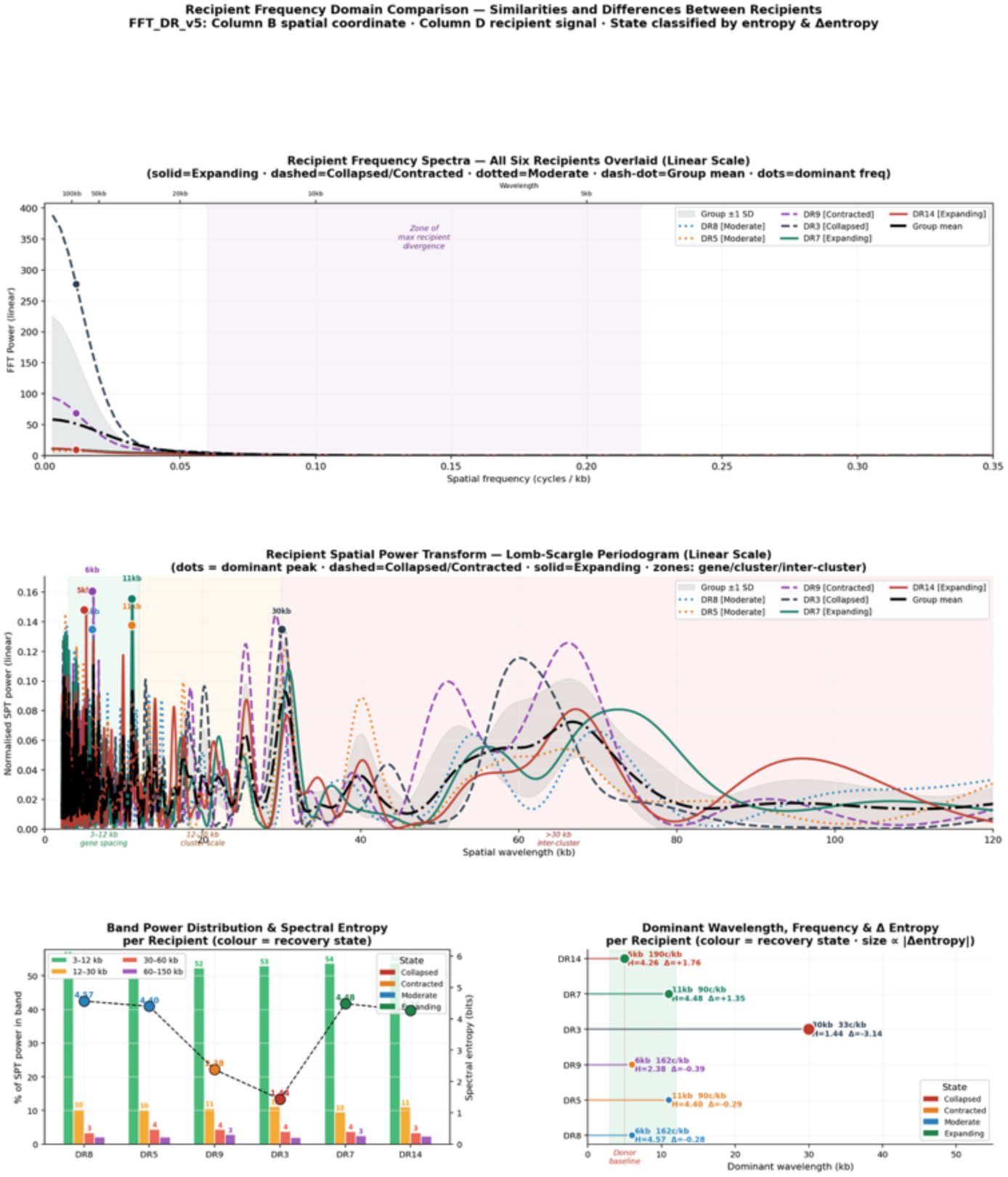

### Donor – recipient comparison

Every donor in every dataset shows its dominant spectral power concentrated at spatial wavelengths of 4–6 kb, corresponding to spatial frequencies of approximately 162–252 millicycles/kb (Figure 3A). Recipients diverged from this in a direction that depends on their recovery state. Collapsed and contracted recipients (DR3, DR9) shifted their dominant wavelength to 6–30 kb (33–162 millicycles/kb), with DR3 showing the most extreme displacement to 30 kb; some recipients (DR7, DR14, DR8, DR5) retained dominant wavelengths in the 5–11 kb range but with altered power distribution above and below that peak. In the spatial power transform (Figure 3D), all donors concentrate 50–56% of their spectral energy in the 3–12 kb sub-family spacing band, with the remaining power distributed across longer wavelengths: approximately 9–10% in the 12– 30 kb cluster band, 3–5% in the 30–60 kb inter-cluster band, and less than 3% beyond 60 kb. This distribution is tightly conserved across donors, with a maximum spread of approximately 6 percentage points across any band. However, recipients show a spread of ∼26% in the 3–12 kb band (28–54%), indicating that some recipients have lost the majority of their short- wavelength power. The 30–60 kb band ranges from 3% (DR8) to 15% (DR3) across recipients, a fivefold range compared to the twofold range seen in donors. When the group-mean recipient FFT spectrum is divided by the group-mean donor spectrum — the power ratio — the pattern is consistent across all analyses. Recipients carry approximately 2–3× more power than donors at wavelengths of 3–5 kb (the fine intra-sub-family scale) and at 20–40 kb (the inter-cluster scale). Donors carry more power than recipients at the intermediate 8–15 kb band, which corresponds to the spacing between different V-gene sub-family clusters. The lower frequency shifts are corroborated by a negative shift in the spectral centroid (SC) in DR3, 5, 8, and 9 (Table 1B), where low SC are associated with lower frequencies, higher low frequency index and lower spectral entropy. Two DRP, 7 and 14 did not follow this trend, suggesting that these recipients were in a more advanced phase of immune recovery than the other DRP analyzed.

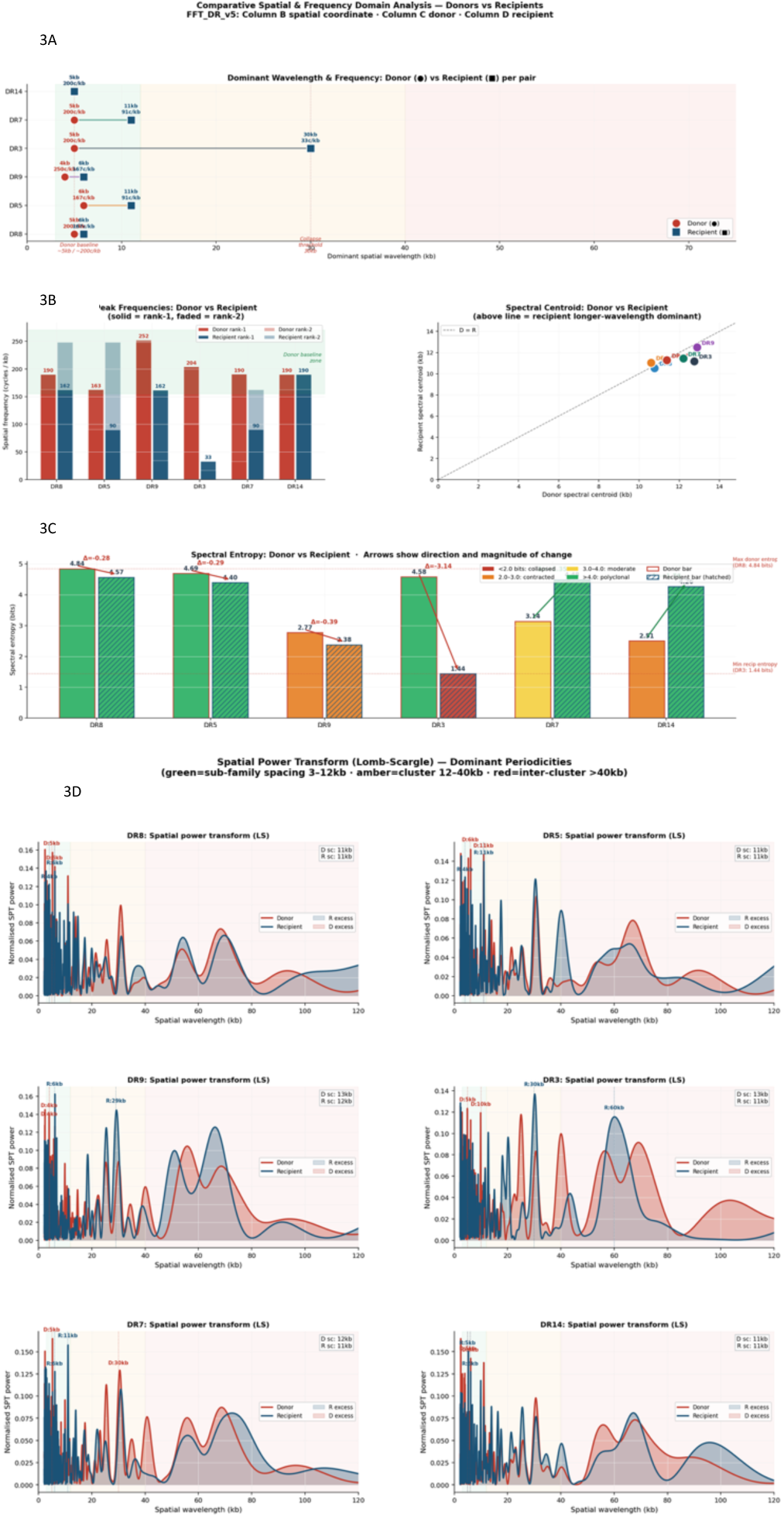

